# Small subunits MttS and MttQ of the MttP transporter regulate trimethylamine transport in *Methanosarcina mazei*

**DOI:** 10.64898/2025.12.02.691841

**Authors:** Tim Habenicht, Lisa Hellwig, Claudia Kiessling, Felix J. Elling, Ruth A. Schmitz

## Abstract

Small proteins (<100 aa) have moved in the focus of science after being overlooked for decades due to bioinformatical and biochemical challenges. While mass spectrometry-coupled ribosome profiling of the mesophilic methanoarchaeon *Methanosarcina mazei* has recently unveiled a wealth of novel small ORFs, the functional roles of most of their products remain unknown. Here, we report the characterization of MttQ (98 aa) and MttS (49 aa), products of small ORFs situated in an operon alongside genes encoding a drug-metabolite-efflux (DME) family transporter (*mttP*) and other enzymes involved in trimethylamine (TMA) degradation. MttS and MttQ interact with MttP to form a stable oligomeric complex spanning the cytoplasmic membrane. TMA transport activity of the MttQ/MttS/MttP-complex is demonstrated via *in vivo Escherichia coli* cells heterologously expressing this system. Based on the reduced growth of a *M. mazei* mutant lacking *mttS*, on TMA as sole carbon source, we conclude that MttS governs the specificity of TMA transport. We posit that interactions between a DME transporter, *e.g.,* MttP, and small proteins fueled evolution of the MttPQS complex and the advent of selective TMA uptake in methylotrophic methanoarchaea. These findings suggest an evolutionary mechanism on how small accessory proteins can alter conserved core functions in order to explore new ecological niches. Further, given that TMA levels in the human bloodstream influence cardiovascular disease risk but can be degraded by host-associated methanoarchaea containing a homolog of the MttPQS-complex, our findings present insight into an archaeal pathway with relevance to human health.

## Introduction

Over the past decade, small open reading frames (ORFs) and their encoded small proteins of 100 amino acids (aa) or less have received increasing attention across various fields of research. This has been driven by advancements in omics technologies, including genome-wide differential RNA sequencing and improved mass spectrometry methods for small proteins, which have illuminated the previously overlooked small proteomes of several prokaryotes (Cassidy et al., 2019; Hadjeras et al., 2023; Kaulich et al., 2021). Ribosome profiling, a powerful technique for mapping actively translated mRNA regions, has proven essential for identifying small ORF-encoded proteins (Brar & Weissman, 2015; Ingolia, 2014). In combination with translation initiation site (TIS) profiling, ribosome profiling promotes the discovery of novel genes (Stringer et al., 2022; Weaver et al., 2019). In addition to facilitating the identification of individual small ORFs, these innovative techniques reveal nested genes within larger ORFs, alternative start sites, and small ORFs embedded within annotated small RNAs (*i.e.*, dual-function RNAs).

Bioinformatic approaches have evolved to overcome biases favoring the detection of large ORFs (> 100 aa) (Orr et al., 2020). Machine learning-assisted tools now offer high-accuracy predictions of small ORFs, enhancing our ability to detect and characterize these elusive yet functionally important proteins. It has recently become increasingly clear that small proteins contribute significantly to a diverse spectrum of cellular processes, including transport and enzyme regulation (Burton et al., 2024; Gray et al., 2022; Hemm et al., 2020; Jordan et al., 2023). Several small proteins identified by biochemical techniques have been shown to serve essential roles in cellular functions, such as ribosomal proteins as small as 38 aa (Nikolay et al., 2015). Unfortunately, however, most novel small proteins remain uncharacterized. Due to their lack of known domains and homologs of established function, automated functional annotation remains extremely challenging, relying heavily on labor-intensive experimental approaches (Fuchs & Engelmann, 2023).

Using ribosome footprinting, 314 novel small ORFs were identified with high confidence in the mesophilic methanoarchaeon *Methanosarcina mazei.* Of these, 26 were verified by LC-MS/MS and 13 by epitope tagging followed by western blot analyses (Tufail et al., 2024). One of the newly identified small ORFs verified by LC-MS/MS and immunoblotting, sORF05, encodes a 49 aa protein. This gene is organized in a predicted operon, situated directly downstream of MM_1691 (*mttP*) and MM_1692 (Deppenmeier et al., 2002; Krätzer et al., 2009). MttP, a membrane protein, belongs to a family of drug metabolite efflux (DME) transporters found ubiquitously throughout the tree of life (Jack et al., 2001). MM_1692 encodes a hypothetical small protein (97 aa) lacking functional annotation (Krätzer et al., 2009; Paul et al., 2000; Tufail et al., 2024).

Most proteins encoded by genes flanking this predicted operon are involved in trimethylamine (TMA) and/or dimethylamine degradation, *e.g.*, TMA methyltransferase MttB (MM_1688), dimethylamine methyltransferase MtbB (MM_1693), and the matching methyltransferase corrinoid proteins MttC (MM_1690) and MtbC (MM_1687) respectively. This entire gene cluster, inclusive of the one predicted operon, is regulated transcriptionally based on TMA availability (Krätzer et al., 2009; Paul et al., 2000; Veit et al., 2005). Compared to their growth on methanol, *M. mazei* cultures grown with TMA as the sole carbon and energy source exhibited 4.93-fold up-regulated transcription of *mttP* (MM_1691), implicating the active involvement of this gene product in TMA metabolism (Krätzer et al., 2009). In *Methanosarcina acetivorans*, the methanoarchaea-specific transcription factor AmzR represses the transcription of the corresponding operon in the absence of TMA. Upon TMA binding, AmzR exhibits decreased DNA-binding affinity to its specific binding box, ultimately resulting in increased transcription of TMA metabolic genes (Ferrer et al., 2025).

We set out to characterize the functions of the products of newly identified sORF05 (*i.e.,* MttS) and MM_1692 (*i.e.*, MttQ) genes, particularly in relation to their neighboring MM_1691 gene predicted to encode a TMA transporter (*mttP*). We deployed an array of bioinformatic, biochemical, and genetic approaches to elucidate the transport activity capabilities, substrate specificity, and interactions of structural and/or functional relevance among these two novel small proteins and MttP.

## Results

### The small proteins, products of two proximal small ORFs, are involved in TMA metabolism

Due to their proximal genomic localization, *M. mazei’s* recently discovered small proteins MttQ and MttS were predicted to be involved in methylamine metabolism. As such, we examined their functional roles and potential involvement in TMA metabolism. The *mttQ* gene (MM_1962) encodes a small protein of 97 aa and is located 1 bp downstream of the *mttP* gene (MM_1691), which encodes a DME family transport protein. Situated 18 bp downstream of *mttQ*’s stop codon is newly identified *mttS*. These three genes are located in close proximity to four other genes involved in full TMA degradation, *i.e.,* TMA methyltransferase MttB (MM_1688) and corresponding methyltransferase corrinoid protein MttC (MM_1690), dimethylamine methyltransferase MtbB (MM_1693), and methyltransferase corrinoid protein MtbC (MM_1687);(Krätzer et al., 2009; Paul et al., 2000; Veit et al., 2006); Fig. 1A). Consequently, these genes were all predicted to be organized in one operon (Krätzer et al., 2009). Reverse transcriptase (RT)-PCR using primer pairs that spanned the end and beginning of each neighboring pair of genes, and RNA extracted from exponentially growing *M. mazei* on TMA confirmed that genes MM_1687 through MM_1693, including *mttS*, are expressed as one single transcript (Fig. 1A, Fig. S1).

**Fig. 1:**
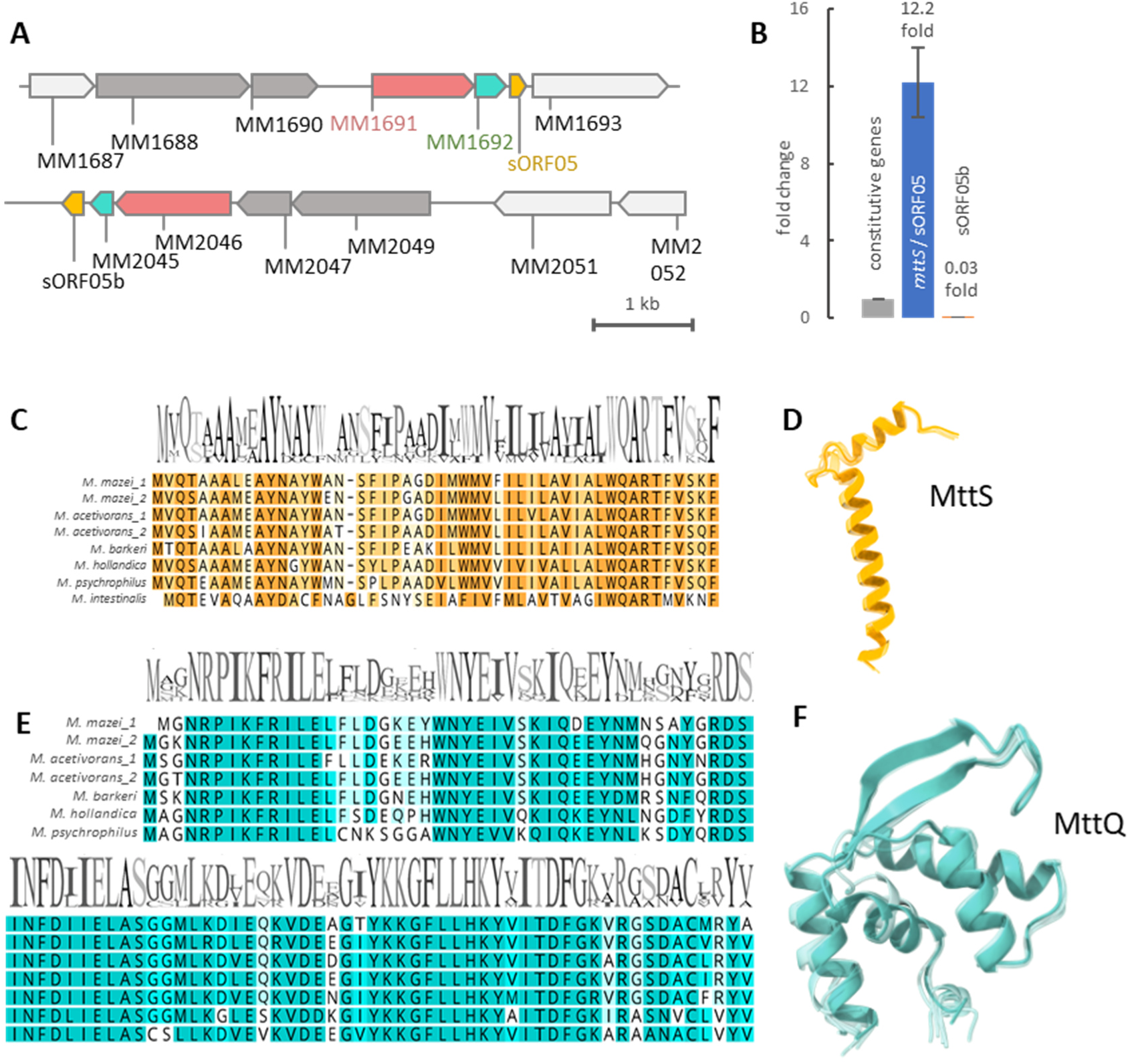
Genomic localization and conservation of MttPQS. **A:** Cartoon of the operon encoding MttP, MttQ, MttS and four flanking genes, all involved in TMA metabolism (MM_1687, MM_1688, MM_1690, MM_1693). The paralogous region (sORF05b, MM_2045 – MM_2052) shown in the lower panel encodes the same genes, with homologous *mttPQS* genes in the same order, but flanking genes in different order. **B:** The transcript level of mttS and the homolog gene (MM_RS07405 = sORF05b) were analyzed by qRT-PCR in comparison with the expression of two constitutively expressed genes (MM_1215 and MM_1621) depicted in grey (1.0 by definition). **C:** The proteins encoded by *mttS* in various methylotrophic archaea exhibit highly conserved aa sequence variability. **D:** MttS ribbon structure predicted by AlphaFold 3 (Abramson et al., 2024) and overlay of predicted structures of the second MttS homolog in *M. mazei* (sORF05b); as well as homologous proteins in other methylotrophic methanoarchaea. **E:** The proteins encoded by *mttQ* genes in various methylotrophic archaea exhibit highly conserved aa sequence variability. **F:** Overlay of predicted structures of MttQ, the paralogue small open reading frame in *M. mazei,* and homolog proteins encoded in several methylotrophic archaea. The homologs of MttQ are structurally highly conserved, based on AlphaFold 3 predictions. *M. intestinalis*, *Methanomassilicoccus intestinalis*; *M. psychrophilus, Methanolobus*; *M. barkeri*, *Methanosarcina barkeri; M. acetivorans*, *Methanosarcina acetivorans*; and *M. hollandica, Methanomethylovorans hollandica*.

The genome of *M. mazei* contains another operon that encodes paralogs of the MttPQS (MM2046/2045, sORF05b) operon. It shows the same suite of genes arranged in different order (Fig. 1A, lower panel, the color code indicates the homologs). Ribosome profiling by Tufail *et al*. and genome wide expression profiling by Krätzer *et al*. showed that this paralogous operon is not transcribed (Krätzer et al., 2009; Tufail et al., 2024), which is corroborated by QRT-PCR analysis. In *M. mazei* cultures grown exclusively on TMA, *mttS* is transcribed in great number (12.2-fold greater than the constitutive genes) while the homologous gene (sORF05b) is barely transcribed at all (0.03-fold compared to the constitutive genes; Fig. 1B).

MttS is highly conserved in Methanosarcinales, including *Methanosarcina* spp. [*e.g*., *M. bakeri* (84 % aa similarity), *M. acetivorans* (94 % aa similarity)], *Methanomethylovorans* spp. [*e.g*., *M. hollandica* (76 % aa similarity)], *Methanolobus* spp. [*e.g*., *M. psychrophilus* (73 % aa similarity)], *Methanomicrococcus* spp. [*e.g*., *M. stummii* (57.1 % aa similarity)], and *Methanolapillus* spp. [*e.g., M. africanus* (55.1 % aa similarity)]. In addition, MttS homologs (41 % aa similarity) are found in Candidatus *Methanomassiliicoccus intestinalis* isolate MGYG-HGUT-02160 (accession NCBI: LR698974.1) from the human microbiome. Highly conserved in aa sequence and predicted structure (Fig. 1C, D), all homologs of MttS have been discovered in methanoarchaeal genomes. They are encoded immediately downstream of a *mttP* homolog and a small ORF that encodes a hypothetical small protein homologous to MM_1692 (MttQ). Much like the case for MmttS, MttQ homologs exhibit highly conserved aa sequence and secondary structure variation in methylotrophic archaea (Fig 1E, F). Only in Candidatus *Methanomassilicoccus intestinalis,* the DME family transporter MttP and the MttQ homolog appear to be fused to one another forming one large protein (585 aa), strongly arguing for a conserved functional relationship between MttP and MttQ.

### AlphaFold 3 predicts a stable complex embedded in the cytoplasmic membrane

AlphaFold 3 (Abramson et al., 2024; Jumper et al., 2021) predicts the structure of a complex formed by MttP, MttQ, and newly identified MttS with high confidence (Fig. 2). According to the statistical parameters derived from the predicted local distance difference test (pLDDT), predicted aligned error (PAE), and predicted template modelling score (ptm = 0.81), the residues are correctly positioned in their local environment and the resulting structure shows a high confidence to match the actual complex in *vivo* (Fig. 2B). Based on the prediction, MttS forms one hydrophobic transmembrane helix and another short helix facing the exterior of the cell, which are linked by a short unstructured loop. MttP consists of ten transmembrane helices that form one large transmembrane unit, and small cytoplasmic and extracellular hydrophilic domains (Fig. 2D). Both its N- and C-terminus reside in the cytoplasm. MttP has one negatively charged side, which likely faces the exterior of the cell (in accordance with the negative outside rule). The surface facing the cytoplasm exhibits negatively charged fractions in addition to the expected positive cytoplasmic surface (Fig. 2C).

**Fig. 2:**
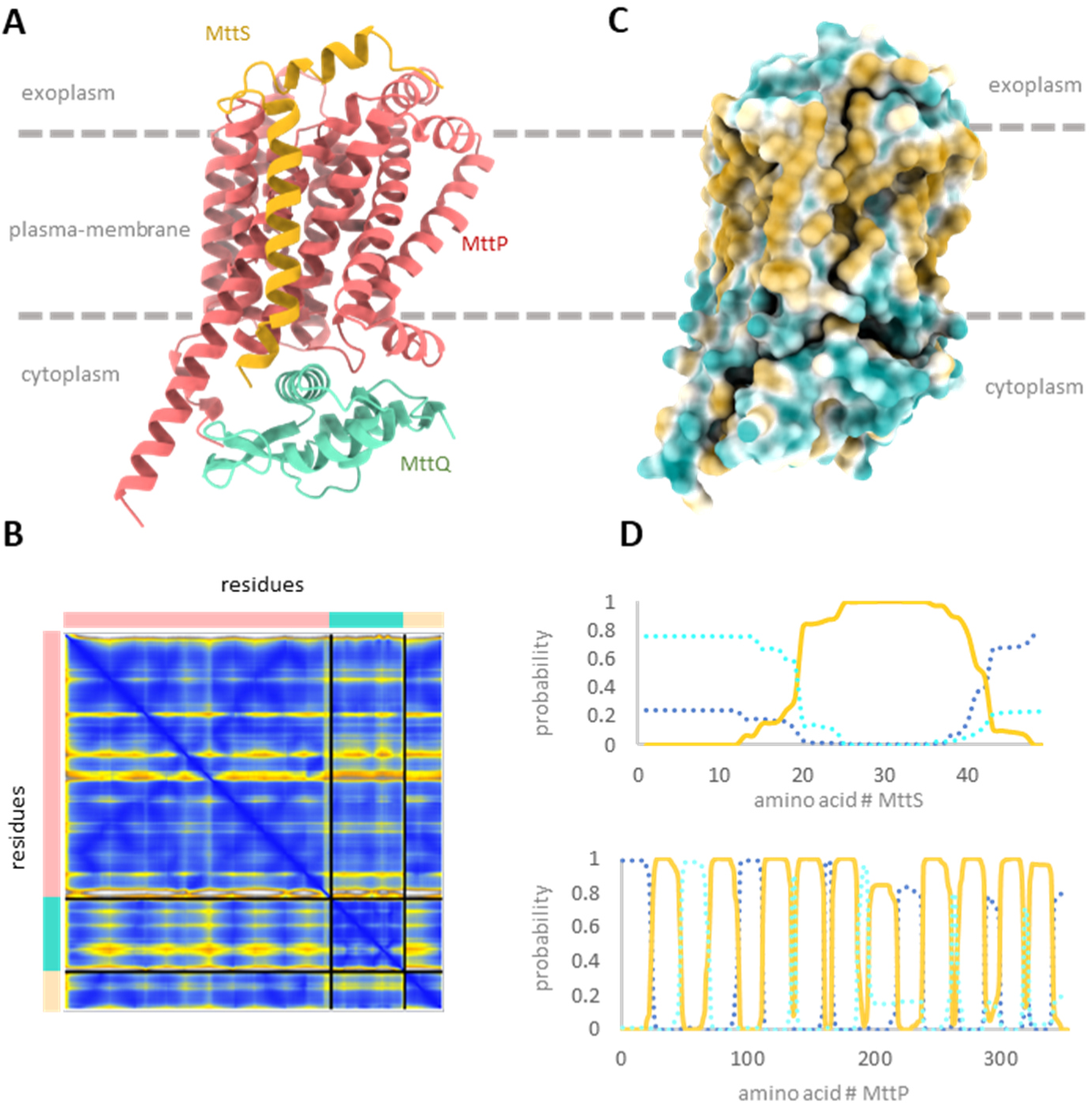
Predicted structure of the MttPQS complex. **A:** AlphaFold 3 prediction (Abramson et al., 2024) of the Mtt-complex consisting of MttP (MM_1691), MttS, and MttQ (MM1692). **B**: Predicted local distance difference test (plDDT) of the Mtt-complex prediction exhibits high confidence in the computational model (dark blue: very high, plDDT > 90; light blue: confident, 90 > plDDT > 70; yellow: low confidence, 70 > plDDT > 50; orange: very low confidence, plDDT < 50). **C:** Hydrophobicity of the surface of the predicted MttPQS complex confirms the likely positioning spanning the cytoplasmic membrane (light blue: hydrophilic; yellow: hydrophobic). **D:** Prediction of hydrophobic transmembrane domains of MttS and MttP by deepTMHMM (Hallgren et al., 2022). MttS consists of one transmembrane helix and a short extracellular domain. MttP envelops ten transmembrane helices, characteristic of DME family proteins (Jack et al., 2001).

MttQ is mostly hydrophilic, consisting of four short helices and two beta sheets. This protein’s surface is divided into one positively charged and one uncharged side. MttQ binds to the membrane embedded MttP/S complex on its cytoplasmic side and is predicted to interact with MttS and MttP in a 1:1:1 ratio (with high confidence; iptm: 0.8; ptm: 0.81; PAE shown in Fig. 2B). The transmembrane helix of MttS is predicted to bind parallel with those of MttP, with a hydrophilic helix of ten aa extending into the extracellular space. In contrast, the hydrophilic MttQ is predicted to bind on the cytoplasmic surface of MttP, whereby the positively charged side of the former covers the negatively charged fraction of the latter’s cytoplasmic surface.

### MttP forms a stable complex with MttQ and MttS

A tag-less version of MttS was generated by purifying His_6_-SUMO-MttS fusion protein expressed in *E. coli* (pRS2106), followed by SUMO protease cleavage (Fig. 3A), which was used to generate antibodies. In parallel, MttP was heterologously expressed with MttQ and MttS in *E. coli* C43 (containing the pRIL plasmid). The expression plasmid (pRS1956) housed the original sequence and organization of all three Mtt genes under the control of the T7-promotor, with the large subunit (MttP= MM_1691) fused to an N-terminal His_5_-tag. His_5_-tagged MttP was purified by Ni-NTA affinity chromatography from solubilized membrane fractions (1 % LMNG). Purified proteins were detected and analyzed via SDS-PAGE, which clearly demonstrated the successful co-purification of all three Mtt proteins, strongly indicative of a stable MttPQS complex.

**Fig. 3:**
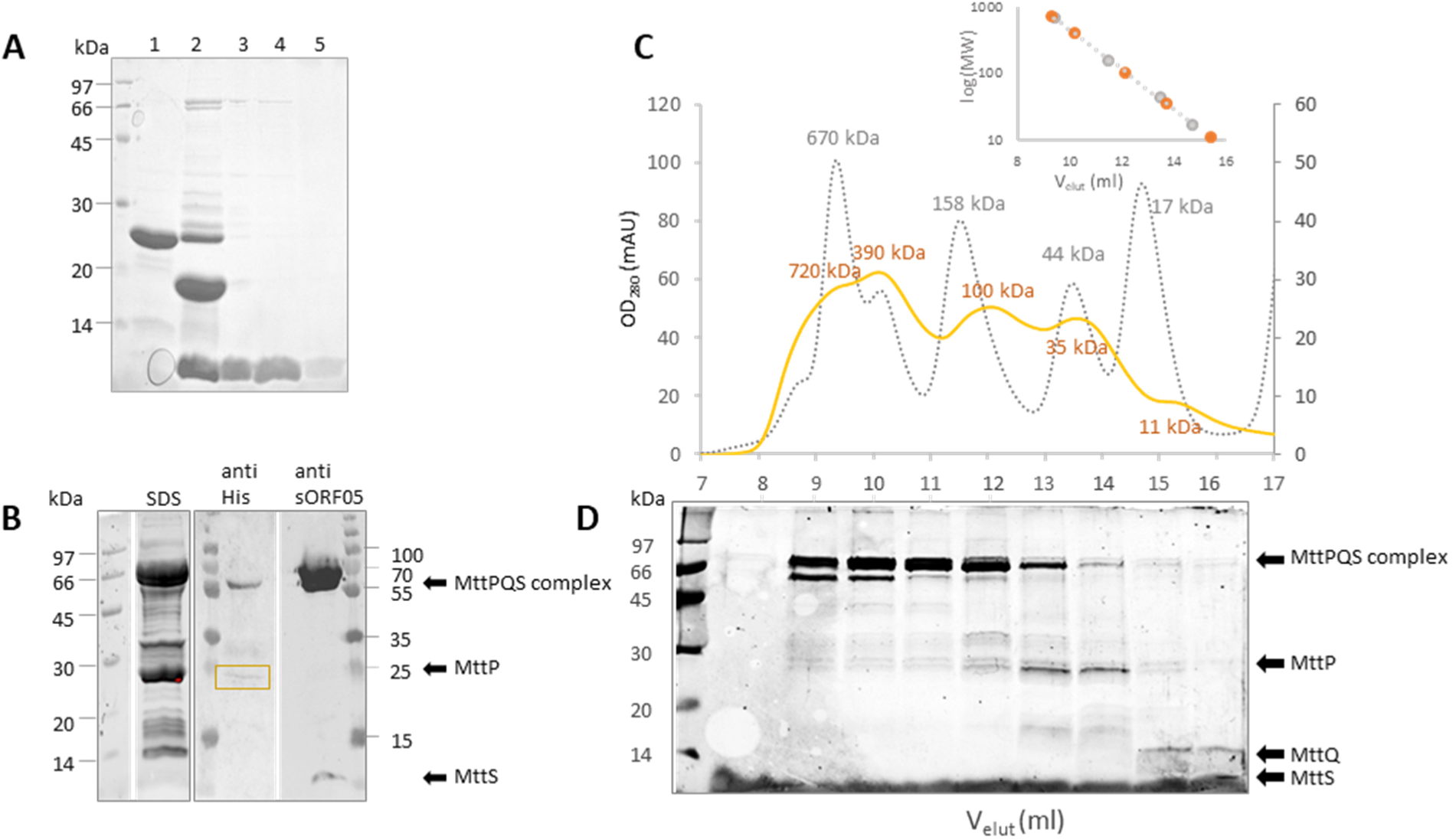
Biochemical characterization of the Mtt complex expressed in *E. coli*. **A:** Purification of heterologously expressed MttS (including cleavage of the SUMO-tag) used for antibody generation. Lane 1, His_6_-SUMO-MttS prior to protease treatment; lane 2, MttS following protease treatment; lanes 3 - 5, MttS following protease treatment and subsequent IMAC purification. **B:** Purification of the Mtt-complex, consisting of His_5_-MttP, MttS, and MttQ (MM1692), from heterologous expression in *E. coli.* Pictured are the results of SDS-PAGE and western blot analyses using anti-His antibodies and those specifically targeting MttS. **C:** SEC chromatogram of the Mtt-complex on Enrich SEC column 650 (BioRad, Hercules, CA, USA). ● (orange): MttPQS complexes, ● (dotted grey): size exclusion standard (BioRad). Peak maxima (elution volume in ml) are plotted against molecular weight [log(MW)] to yield the calculated molecular weight of the eluted proteins (insert). D: Elution fractions resulting from SEC on SDS-PAGE show the presence of MttP, MttS, and MttQ in different peaks/complexes. For molecular masses, see also supplemental Table S1.

SDS-PAGE analyses revealed bands at roughly 65, 55, 35, 25, 12, and 6 kDa, most likely representing the three expressed proteins in various combinations (Fig. 3B). Western blot analyses exploiting antibodies against the His-tag detected His_5_-MttP in the 25 kDa signal, corresponding to monomeric MttP. While this is below MttP’s predicted molecular weight of 35 kDa, this is a common phenomenon associated with membrane proteins, which often migrate faster than expected on SDS-PAGE due to incomplete denaturation and atypical detergent interactions (Rath et al., 2009). In addition to the monomer, His_5_-MttP was also detected in the 55 kDa signal (Fig. 3B, mid panel), hinting at an SDS-stable MttPQS complex. When a specific antibody directed against MttS was used to evaluate the same elution fraction on a separate blot, results clearly demonstrated the presence of this small protein in both 6 kDa monomeric and 55 kDa complex-associated forms (Fig. 3B, right panel). These results show the presence of, and thus stable interactions between, MttS and MttP in the 55 kDa protein complex, even in the presence of SDS. While the third subunit, MttQ, could not be monitored directly, the molecular mass of the 55 kDa complex is consistent with 1:1:1 stoichiometry of all three subunits.

To verify the formation of subunit complexes, purified elution fractions were subjected to size exclusion chromatography (SEC). Resulting chromatograms revealed five distinct peaks (Fig. 3C), corresponding to different oligomeric states, combinations of subunits, and individual subunits. Elution volumes coincided with approximate molecular masses of 720, 390, 100, 35, and 11 kDa. These results indicate that MttP, MttQ, and MttS can form higher-order oligomeric complexes. Specifically, the 100 kDa peak matches the expected size of a dimer of the ∼55 kDa heterotrimeric complex comprising MttP, MttQ, and MttS. The three largest peaks likely represent complexes consisting of two (100 kDa), seven (390 kDa), and thirteen (720 kDa) protomers (Table S1). SDS-PAGE analyses of these SEC fractions confirmed the presence of MttP, MttQ, and MttS, supporting the notion that they assemble into stable, high-molecular-weight oligomers in a 1:1:1 ratio (Fig. 3D).

The 35 kDa peak observed via SEC likely represents monomeric MttP in the absence of interaction partners, as evidenced by a 25 kDa signal via both SDS-PAGE and western blot analyses, which is consistent with prior MttP purifications. The 11 kDa peak corresponds to monomeric MttQ (MM_1692). Due to its small size, MttS (∼ 5 kDa) might not be separated adequately via the SEC approach. Regardless, SDS-PAGE analysis of the smallest peak confirmed the presence of a corresponding signal.

### **The MttPQS complex actively transports TMA** *in vivo*

*E. coli* does not encode a TMA transporter and cannot degrade TMA. As such, the *M. mazei* Mtt complex was heterologously expressed in *E. coli* from plasmid pRS1956, and *E. coli* cultures carrying the empty plasmid pET28a served as a null control. Cultures growing exponentially in nutrient rich Luria Bertani (LB) medium were supplemented and incubated with TMA (33 mM) for one hour, harvested, washed with PBS and lysed in deionized water. Cleared lysates were evaluated via high performance liquid chromatography (HPLC) coupled to mass spectrometry (MS), to determine TMA content. TMA standards ranging in concentration from 1 to 10 µg TMA/ml were used for calibration. When the empty vector control was analyzed, only low concentrations of TMA were detected most likely due to unspecific transport or insufficient washing. The lysate derived from cells expressing the Mtt complex exhibited a 3.5-fold increase in TMA (Fig. 4A). This increase in intracellular TMA concentration in the mutant (and not the control strain) was observed in two independent biological replicates, clearly demonstrating the TMA transport activity of the Mtt complex. In addition, these results show that the Mtt complex is active completely independent of proteins of the flanking genes, even in the heterologous *E. coli* environment.

**Fig. 4:**
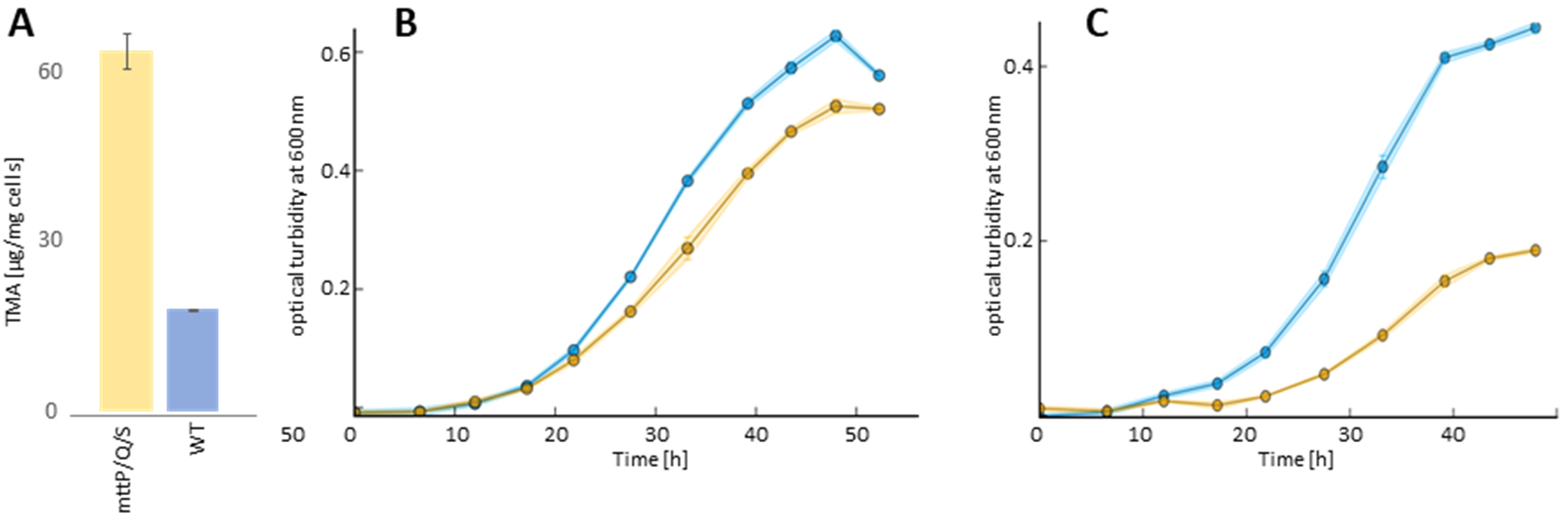
MttS is required for efficient transport of TMA. **A:** TMA transport assay with *E. coli* cells expressing the MttPQS complex from a plasmid. Heterologously expressed MttPQS-complex enables *E. coli* to import TMA, in contrast to the *E. coli* control strain. TMA was measured by LC-MS/MS. Depicted are the mean values of two biological replicates. The error bars show the standard deviation. **B, C:** Growth of the *M. mazei* Δ*mttS* mutant strain (yellow) and wt (blue) on methanol (B), and TMA (C). For all growth curves one of four biological replicates is shown. For each biological replicate, three technical replicates were measured, of which the mean values are depicted. The error bars and the colored areas show the standard deviations.

### MttS is required for Mtt complex-facilitated TMA transport

To verify MttS involvement in transport activity, the chromosomal gene encoding MttS was replaced with a puromycin resistance cassette resulting in a *M. mazei* Δ*mttS* mutant strain. Growth characteristics were compared between the mutant and wild type strain (wt) supplied with either 30 mM methanol or 10 mM TMA as a sole source of carbon and energy. The wt and Δ*mttS* mutant strain exhibited no significant differences in growth on methanol (Fig. 4B). However, when growing on TMA, wt cells proliferated at the same rate they did on methanol, following a short adaptation phase (likely to induce TMA degradation pathways). In contrast, the absence of small protein MttS resulted in a significantly hindered growth rate and complete cessation of proliferation upon reaching a turbidity of 0.15 (at 600 nm; T_600_) in four independent biological replicates (Fig. 4C). This clearly demonstrates that the MttPQS complex is responsible for TMA transfer and small protein MttS is required for efficient transport activity.

### Predicted TMA transport channel

The AlphaFold3 predicted structure of the MttPQS complex was used for analysis of a potential transport channel. Using the software Mole 2.5 (Berka et al., 2017), a continuous channel along the membrane spanning axis of the protein complex was identified. The total length of the predicted pore was 32.4 Å, spanning from the extracellular to the cytoplasmic side (Fig. 5). However, the pore shows several narrow constriction points with a radius of less than 2.5 Å (Fig. 5B) which do not allow free diffusion of trimethylamine (TMA), whose effective radius is about 2.6–2.8 Å. This indicates that substrate passage likely requires conformational changes during the transport cycle rather than a permanently open channel.

**Fig. 5:**
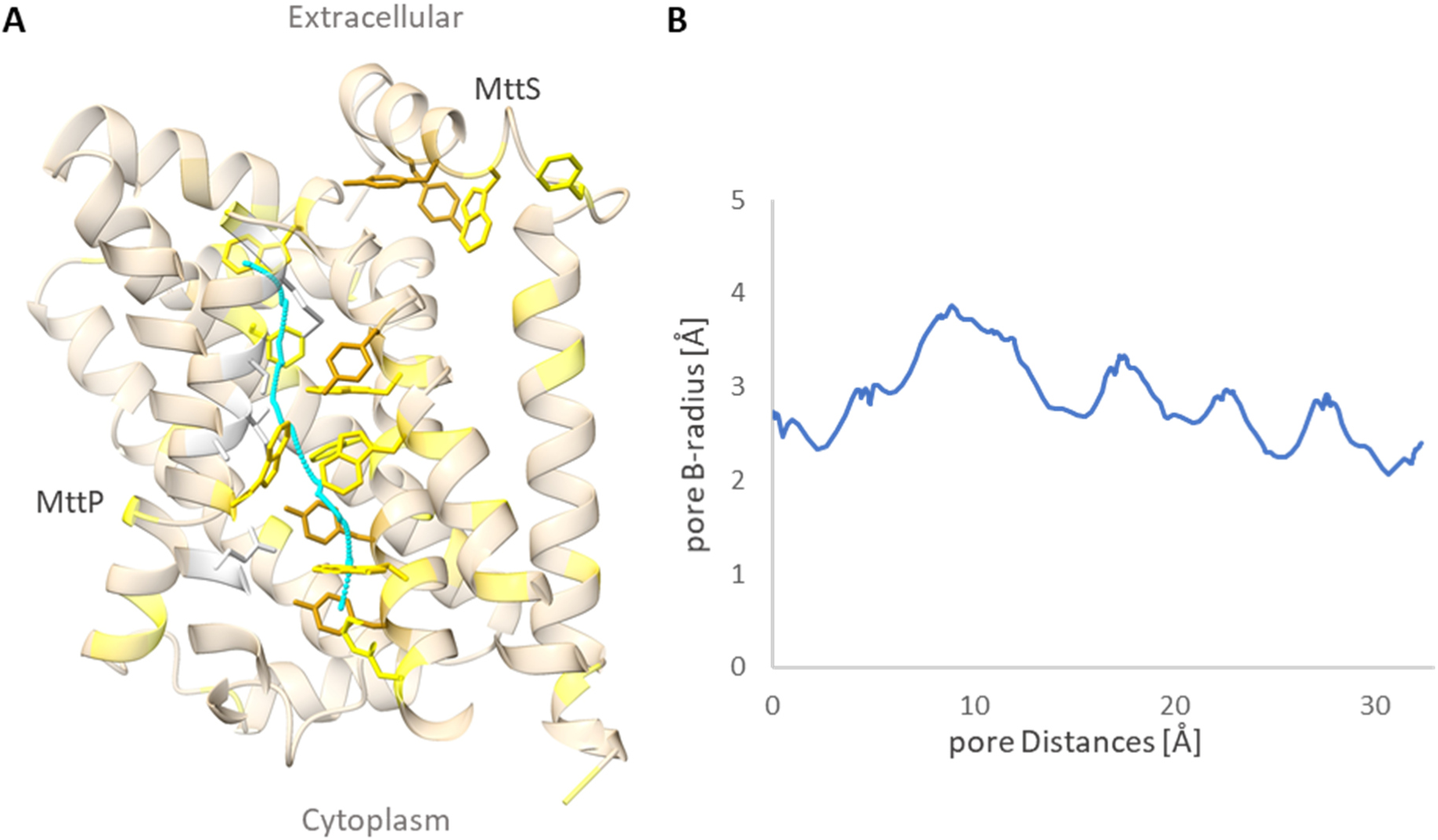
Analysis of the pore spanning MttP. **A:** AlphaFold3 structure of MttPQS showing only MttP and MttS. The Pore identified by MOLE 2.5 analysis is depicted as a cyan line. All residues in a maximal distance 3 Å of the pore are shown including sidechains. Those lining residues consist mostly of aromatic amino acids Trp, Tyr and Phe. In addition, aromatic amino acids in the extracellular domain of MttS are shown with their respective side chains. These residues might be involved in TMA binding. **B:** The pore B-radius (which includes estimated errors of the AlphaFold3 prediction) as analyzed by MOLE 2.5 shows a continuous channel with a mean pore radius of 3 Å.

Further analysis of the structure revealed a high number of aromatic residues (Phe, Tyr, Trp) lining the pore (Fig. 5A), several of which cluster around the narrow regions. These aromatic residues might form a cation–π interaction network, stabilizing the positively charged TMA as it passes through the hydrophobic core.

Similar, the extracellular domain of MttS also contains a relative high number of Trp and Tyr (Y11, Y14 and W15), which are residues potentially involved in extracellular TMA binding Fig. 5a).

## Discussion

We investigated the roles of two previously uncharacterized yet highly conserved small proteins, MttS (49 aa) and MttQ (98 aa), in TMA metabolism, and show that both proteins form a stable complex with the DME family transporter MttP. This tripartite complex mediates TMA uptake. Moreover, the chromosomal deletion of *mttS* significantly impairs the growth of *M. mazei* when TMA is the sole source of carbon and energy.

Recently, the *mttS* gene was identified by ribosome profiling as ribo_sORF05 and confirmed at the protein level via immunoblotting (Tufail et al., 2024). Previously annotated as a conserved hypothetical protein based on sequence homology and only recently verified at the protein level by mass spectrometry (Tufail et al., 2024), MttQ has yet to be functionally characterized. The genes encoding each of these proteins are situated within a putative operon, which we validated experimentally using RT-PCR. This operon, encoding a total of seven proteins (Fig. 1A), is upregulated upon utilizing TMA as a sole carbon and energy source (Krätzer et al., 2009; Veit et al., 2005). In addition to proteins involved in TMA degradation, this operon includes genes for dimethylamine (DMA) degradation. The first gene in the operon (MM_1687) encodes the DMA corrinoid-binding protein, while the last gene (MM_1693) encodes the methyltransferase responsible for transferring a methyl group from DMA to coenzyme M (CoM), thereby generating monomethylamine (Ferguson et al., 2000). This arrangement may reflect evolutionary operon expansion via the recruitment of genes involved in downstream pathways (degrading DMA, the product of the first step), thereby enhancing fitness by enabling synchronized expression of TMA and DMA degrading proteins.

The MttPQS complex was successfully expressed from a plasmid as one artificial operon in *E. coli*, and co-purification experiments using tagged MttP demonstrate that MttS and MttQ physically interact with MttP to form a stable protein complex (Fig. 3). The formation of this complex aligns with computational predictions (Fig. 2), and structural modeling indicates that MttP comprises ten transmembrane helices and a small cytoplasmic domain, all of which are characteristic of DME transporters. MttP is distinguished from other DME family proteins by its strong and specific interactions with MttS and MttQ. These small proteins are not present in any other bacteria or (methano)archaea, except methylotrophic methanogens. MttS is predicted to bear a single transmembrane helix and a hydrophilic extracellular domain, whereas MttQ (MM_1692) is a soluble protein composed of α-helices and β-sheets, likely associating with the cytoplasmic face of MttP. SEC analyses suggest that the Mtt complex forms higher-order oligomers, potentially consisting of 7 to 14 protomers in 1:1:1 stoichiometry (Fig. 3C, D).

Based on its genomic co-localization with *mttB* (a TMA methyltransferase) and *mttC* (the associated corrinoid protein), and its organization in one operon, MttP was hypothesized to encode a TMA-specific transporter (Krätzer et al., 2009). However, this hypothesis was never confirmed empirically. By overexpressing the MttPQS complex in *E. coli*, we successfully engineered a TMA-transporting bacterium, thereby providing the first direct experimental evidence of MttP’s TMA-transport function (Fig. 4A).

To date, the only other known TMA transporter is TmaT from the deep-sea bacterium *Myroides profundi* (see Fig. 6a), which belongs to the betaine-choline-carnitine-transporter (BCCT) family (Ziegler et al., 2010). Unlike *M. mazei*, *M. profundi* does not catabolize TMA but converts it to trimethylamine N-oxide (TMAO), which serves as a compatible solute for osmotic and pressure stress adaptation (Qin et al., 2021). The structure of TmaT was recently reported, which gives first insights in the molecular mechanisms behind TMA transport. The cryo-EM structures of TmaT binding TMA show aromatic amino acids Trp and Tyr to bind and stabilize TMA during transport (Gao et al., 2025).

**Fig. 6:**
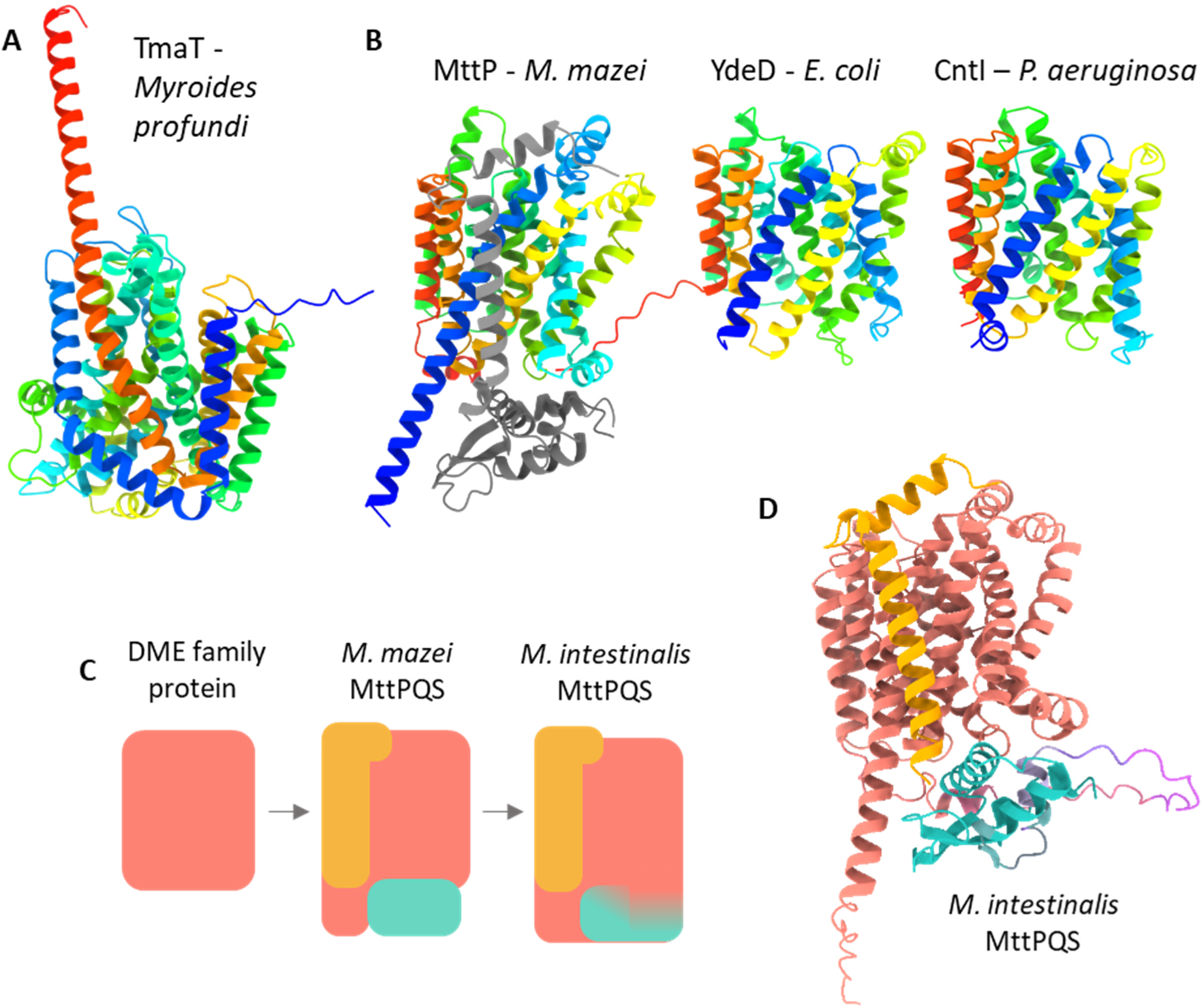
Evolution of transport protein structures. **A+B:** Structural predictions of the TMA transporter TmaT and three DME family transporters. **A:** TmaT of *M. profundi*. **B:** MttP of *M. mazei;* YdeD of *E. coli*; and Cntl of *Pseudomonas aeruginosa*. All Proteins are colored by rainbow from N-terminus (red) to C-terminus (blue). Small proteins MttS and MttQ are added in grey to MttP. **C:** Proposed evolution of the MttPQS complex. The evolutionary old DME family transporter evolved to be an TMA importer by interaction with newly evolved small proteins MttQ and MttS. In *M. intestinalis* MttQ is fused to the main protein as a further adaptation. **D:** AlphaFold3 predicted structure from the *M. intestinalis* MttPQS complex. MttP (red) is linked to MttQ (cyan) by a linker sequence (purple). MttS is depicted in orange.

Analysis of the predicted MttPQS structure revealed a continues pore running through the predicted transporter MttP of *M. mazei*. Along the membrane spanning axis, a solvent-accessible pathway is proposed that extends from the cytoplasmic to the extracellular side (Fig. 5A). A profile of local pore radius along this axis showed numerous narrow constrictions (Fig. 5B). The minimal estimated radius was approximately 2.5 Å, indicating a narrow gate that would require conformational flexibility for substrate translocation. Besides the fact that the structure is a static AlphaFold prediction and has to be analyzed as such with reasonable care, such constrictions are consistent with transporters that undergo alternating-access motions rather than acting as an open channel (Diallinas, 2014; Kaback & Smirnova, 2011). The lining is dominated by aromatic residues, interspersed with few hydrophobic or negatively charged aa, mainly Trp and Tyr similar to the proposed pocket in the TmaT from *M. profundi* (Gao et al., 2025). The presence of aromatic rings at the constrictions suggests the potential for cation–π interactions, while nearby acidic residues may provide complementary electrostatic stabilization. Their presence suggests a selectivity mechanism favoring small amine-like substrates, consistent with a role in TMA recognition and translocation. Consequently, the predicted structural features support a model where TMA transport occurs via a gated, substrate-specific mechanism, with aromatic residues forming a transiently permissive pathway rather than a permanently open pore. Taken together, the geometric continuity of the pore, the size and distribution of the constriction, and the enrichment of aromatic and acidic residues along the channel wall all support the interpretation that MttP functions as a TMA transporter.

DME family transporters are found throughout all domains of life and are generally implicated in the export of metabolic byproducts and toxic compounds (*e.g*., YdeD of *E. coli,* CntI of *Pseudomonas* spp.) (Daûler et al., 2000; Gomez et al., 2021; Lhospice et al., 2017). However, functionally characterized DME proteins are scarce, and hitherto no experimental structures have been resolved. High-confidence computational models imply conserved architecture across the family despite low sequence similarity (Fig. 6B). MttS and MttQ are conserved exclusively in methylotrophic methanoarchaea, *e.g.*, *Methanosarcina* spp., which utilize a diverse array of carbon sources in addition to CO₂, such as methanol, TMA, and other methylated compounds. This restricted conservation pattern suggests that MttS and MttQ evolved specifically to confer TMA specificity/selectivity to MttP in methylotrophic methanoarchaea. This hypothesis is strongly supported by the growth characteristics of a chromosomal *mttS* deletion mutant of *M. mazei*. When grown on TMA as the sole carbon and energy source, the mutant exhibits a severely inhibited growth rate and reaches a significantly lower final cell density than its wild type counterpart (Fig. 4C). This growth phenotype argues for impaired TMA uptake, especially under conditions of decreased extracellular TMA availability. Based on these findings, we posit that MttS, via its extracellular domain, plays a key role in TMA recognition and substrate specificity of the transporter (especially under TMA-limiting conditions). One attractive hypothesis is that MttS, using its extracellular extension, acts as a sensor or pre-binding lure that increases TMA uptake efficiency under TMA-limiting conditions, functioning analogously to periplasmic-binding proteins in ABC transporters (Saurin et al., 1989). In addition, the presence of Trp and Tyr residues in the MttS extracellular domain is strengthening the proposed function of MttS as a TMA binding domain which increases the TMA sensitivity of the transporter. As a cytoplasmic soluble protein, MttQ, likely also contributes to substrate specificity and/or regulation, though its precise function remains elusive.

Beyond their ecological role, TMA and its oxidized form TMAO have clinical relevance: both are linked to human cardiovascular diseases such as atherosclerosis (Roncal et al., 2019; Zeisel & Warrier, 2017). TMA is produced by gut bacteria depending on the diet (Zeisel & Warrier, 2017; Zhang et al., 1999) and consequently, efficient TMA uptake and degradation by methanoarchaea with as little retention in the intestine as possible helps prevent absorption by the human host and thereby lower the risk of cardiovascular diseases. Methylotrophic methanoarchaea have been reported as part of the human gut microbiome as for instance *Methanomassilicoccus intestinalis* (Nkamga et al., 2017), which encodes a fused MttPQ and a MttS homolog in an operon with TMA and DMA specific methyltransferases (MttBC and MtbBC) (Borrel et al., 2013). Thus, the results of this study not only provide fundamental information on TMA metabolism, but also concrete targets for potential improved TMA uptake in probiotic methanoarchaea. The fact, that TMA import by the MttPQS complex is independent from the host cell, further indicates the potential transferability of the TMA transport into classic probiotic candidates as a direct application.

We conclude that the Mtt complex (*i.e.,* MttP-MttQ-MttS) is a functional TMA importer in *M. mazei*. While MttP shares structural characteristics with other DME transporters, its association with two small proteins appears unique and essential for TMA specificity. Results suggest that, in contrast to the more general efflux roles of other DME transporters, the MttPQS complex represents a specialized adaptation in methylotrophic methanoarchaea, bolstering fitness by facilitating specific and efficient uptake of TMA. In *M. intestinalis*, the fusion of MttP and MttQ (Fig. 6C, D) may represent a further adaptation that allows for more rapid regulation of TMA transport in response to fluctuating environmental TMA concentrations. Taking together, the MttPQS complex might exemplify the evolution of small accessory proteins to confer substrate specificity to a broadly conserved, evolutionarily dated transporter scaffold (Fig. 6B, C).

## Methods

### Construction of plasmids

A cloning plasmid was constructed to facilitate the seamless cloning of SUMO tagged fusion proteins. Bsa1 cleavage sites and mScarlet, a fluorescent marker gene, were cloned into pETSUMO (Qiagen, Hilden, Germany). Plasmid pETSUMO was amplified using specific primers (Eurofins, Ebersberg, Germany, listed in Table S3) while introducing BsaI, BamHI, and NotI cleavage sites. The gene encoding mScarlet, inclusive of both its promotor and terminator sequence, was amplified from pRS1736. Both products were cleaved by the restriction enzymes BamHI and NotI (New England Biolabs, Ipswich, USA) and ligated using T4 ligase (Thermo Fisher Scientific, Waltham, USA). The resulting plasmid was named pRS2100.

The gene encoding MttS was amplified from *M. mazei* Gö1 genomic DNA using specific primers equipped with flanking restriction sites (Eurofins Ebersberg, Germany, Table S3). Both the resulting PCR products and plasmid PRS2100 were digested with BsaI restriction enzymes (New England Biolabs, Ipswich, USA) and ligated with T4 ligase (Thermo Fisher Scientific, Waltham, USA) in a one-step golden gate cloning. The resulting plasmid, bearing the SUMO-MttS fusion under the control of a T7 promoter, was named pRS2106.

To express the MttPQS complex, genes *mttP*, *mttQ*, and *mttS* were cloned into the pET28a expression vector under the control of a T7 promoter, generating a His_5_-MttP fusion and untagged MttQ and MttS. Genes were amplified from chromosomal *M. mazei* DNA using specific primers equipped with flanking restriction sites (Eurofins, Ebersberg, Germany; Table S3), and cloned into pET28a (Table S2) with NdeI, NotI (New England Biolabs, Ipswich, USA), and T4 ligase (Thermo Fisher Scientific, Waltham, USA). The resulting plasmid was named pRS1956 (Table S2).

### Construction of ΔmttS

The 1-kb genomic regions flanking *mttS* on both sides were amplified from *M. mazei* Gö1 genomic DNA using specific primers (Eurofins, Ebersberg, Germany) listed in Table S3. Plasmid pMCL210 (Nakano et al., 1995) was used as a suicide vector for allelic replacement in *M. mazei* (Hüttermann & Schmitz, 2024). The 1-kb upstream and downstream fragments were inserted into plasmid pMCL210 with restriction enzymes *Bam*HI, *Sac*I and *Xho*I. A puromycin resistance cassette (pac; (Metcalf et al., 1997), encoding the puromycin *N*-acetyltransferase gene from *Streptomyces alboniger* under the control of the *M. mazei* constitutive promoter pmcrB, was excised from pRS207 (Table S2) with *Bam*HI and cloned between the two 1-kb flanking regions. The resulting plasmid, designated pRS2228, was linearized using *Sal*I and transformed into *M. mazei* via liposome-mediated transformation (Ehlers et al., 2005). The *mttS* gene was replaced with the *pac*-cassette by simultaneous homologous recombination in both flanking regions (Ehlers et al., 2005). Successfully mutated cells bearing the *mttS* deletion were selected in puromycin-containing medium and plated to obtain single mutant clones (Deppenmeier et al., 2002). Success of the deletion was confirmed via southern blotting using specific probes against the *mttS* gene and the *pac* cassette (Fig. S2).

### Purification of heterologously expressed MttS

*E. coli* BL21 (DE3)/pRIL cultures containing the expression plasmid pRS2106 were continuously shaken in Terrific Broth (TB) at 37°C. At an optical turbidity (T_600_) of 0.8, protein expression was induced by adding 100 µM IPTG (final conc.). Following an additional 3 h incubation, cells were harvested via centrifugation at 4,000 x g at 4°C for 20 min. Cell pellets were resuspended in 6 ml phosphate buffer A (50 mM phosphate, 300 mM NaCl, pH 8.0) and lysed by passing through the French pressure cell twice at 80 N/mm^2^. To remove unlysed cells and cellular debris, the cell extract was centrifuged for at 7,000 x g at 4°C for an additional 30 min.

SUMO-tagged MttS was purified by IMAC using NiNTA on gravity columns with 1 ml bed volume (Cube Biotech, Monheim, Germany). Elutions were conducted using phosphate buffer (50 mM phosphate, 300 mM NaCl, pH 8.0) plus imidazole at 100, 250, and 500 mM concentrations. Afterwards, buffer exchange facilitated removal of the imidazole. Resulting protein fractions were incubated with SUMO protease UlpI (Peroutka III et al., 2011) to cleave all of the His-SUMO tags, at which time the sample was purified by IMAC to remove cleaved tags and all remaining uncleaved fusion proteins. High purity, unlabeled MttS was then sent to Davids Biotechnology (Regensburg, Germany) for anti-MttS-specific antibody production.

### Purification of the heterologously expressed Mtt complex

To purify the Mtt complex, His_5_-MttP, MttS, and MttQ were first expressed from plasmid pRS1956 in *E. coli* C43 pRIL cells (Table 2). Cultures were grown in TB media to T_600_ of 0.8, expression was induced by adding 100 µM IPTG (final concentration), and cells were incubated for an additional 3 h at 37°C. Cultures were then harvested and lysed as described above. Subcellular fractionation of the cell extract was achieved via centrifugation at 200,000 x g at 4°C for 1 h. The supernatant (cytoplasm) was discarded and the pellet was washed in 50 mM Tris/HCl buffer (pH 7.6) and centrifuged again at 200,000 x g at 4°C for 1 h. The resulting pellet, themembrane fraction, was then solubilized by adding LMNG (final concentration 1 %), gently shaken for 1 h at 4°C, and subjected to centrifugation at 200,000 x g and 4°C for 1 h.

His-tagged MttP was purified via IMAC using Ni-NTA on gravity columns with 1 ml bed volume. Elutions were conducted using phosphate buffer (50 mM phosphate, 300 mM NaCl, pH 8.0) plus imidazole in final concentrations of 100, 250, and 500 mM. Following SDS-PAGE, elution fractions in 100 mM imidazole were pooled and dialyzed against phosphate buffer (50 mM phosphate, 300 mM NaCl, pH 8.0) to remove the imidazole.

### SEC analyses

Separation of MttPQS complex subunits was achieved via size exclusion chromatography (SEC) using an Äkta pure chromatography system (Cytiva, Marlborough, USA) outfitted with a gel filtration column Enrich TM SEC 650 (BioRad, Hercules, USA). Proteins were eluted at a flow rate of 1 ml/min in phosphate buffer (50 mM phosphate, 300 mM NaCl, pH 8.0). For every local maximum of the size exclusion chromatogramm, the corresponding molecular mass was calculated by interpolation to the size exclusion standard (Bio-Rad size exclusion standard; #151-1901, Bio-Rad, Hercules, USA).

### **Growth of** *M. mazei*

*M. mazei* was cultivated in either 1 L anaerobic minimal medium in 2 L sealed bottles or in 50 ml anaerobic minimal medium in 80 ml serum bottles under a gas phase consisting of N_2_ and CO_2_ in a 80/20 (vol/vol,) ratio (2 bar). Immediately prior to inoculation, the medium was augmented with 150 mM methanol or 330 mM TMA. *M. mazei* cells were harvested by centrifugation at 6,000 g at 4°C for 45 min, and cells were lysed using a dismembrator (Sartorius, Göttingen, Germany) at 1,600 rpm for 3 min in Tris/HCl buffer (50mM, pH 7.6).

Growth analysis was conducted with the mutant and the wildtype strain (wt). For the growth analysis, cultures with either 30 mM methanol or 10 mM TMA were inoculated with the same preculture grown on methanol.

### RNA extraction and RT-PCR analyses

To extract RNA, 50 ml *M. mazei* cultures were grown as described above to T_600_ of 0.8, at which time they were allowed to cool and then harvested via centrifugation at 4,000 x g at 4 ◦C for 30 min. Cell pellets were resuspended in ROTI®Zol (Carl Roth, Karlsruhe, Germany), and total RNA was isolated via chloroform extraction, followed by DNase I treatment and phenol–chloroform precipitation. Reverse transcriptase polymerase chain reactions (RT-PCR) were performed using the One*Taq*® One-Step RT-PCR Kit (New England Biolabs, Ipswich, USA) and primer pairs designed to bind to two successive genes of the operon. Primer sequences are provided in Table S3. Quantitative reverse transcriptase polymerase chain reactions (QRT-PCR) were performed using the ViiA™ 7 Real-Time PCR System (Applied Biosystems, Darmstadt, Germany) with the PowerUp™ SYBR™ Green Master Mix (Thermo Fisher Scientific, Waltham, MA, USA) and primer pairs designed to bind to *mttS* (MM_RS18880)*, the mttS-homolog (MM_RS18885)* as well as constitutively expressed genes (MM_1216 and MM_1621). Primer sequences are provided in Table S3. Quantitative RT-PCR data were analyzed by comparing ct-values to the housekeeping genes. For each growth condition, three independent biological replicates were conducted.

### Transport assay and HPLC-MS analyses of trimethylamine

The MttPQS complex was heterolougusly expressed in *E. coli* from plasmid pRS1956, as described above. Cells were grown to T_600_ of 0.8, at which time expression was induced by adding 100 µM IPTG (final concenzrtatuon). Following an additional 2 h incubation, to ensure expression of the MttPQS complex, TMA was added (final concentration 33 mM), and cultures were incubated for one hour at 37°C. Cells were then harvested by centrifugation at 6,000 x g, 4°C for 30 min. The resulting pellet was washed twice in phosphate buffer (50 mM phosphate, 300 mM NaCl, pH 8.0), finally resuspended in deionized water (final: 1 ml per 1 mg cell pellet) and lysed by passing through the French Press two times at 80 N/mm^2^. The lysate was centrifuged for 30 min at 13,000 x g. The supernatant was then diluted 1:10 in acetonitrile and subjected to centrifugation at 20,000 x g for 5 min at RT. The resulting supernatant was used for high performance liquid cromatography (HPLC) in water/acetonitrile in a 1:9 ratio. Subsequent mass spectrometry (MS) analyses were implemented to determine TMA content. Known concentrations of TMA, ranging from 1 to 10 µg/ml of TMA in 100 % acetonitrile, were used for calibration.

TMA concentrations in the respective lysates of the transport assay were determined using an Agilent 1260 Infinity II series high performance liquid chromatograph coupled to an Agilent 6130 single quadrupole mass spectrometer (HPLC-MS). TMA and other metabolites were separated via isocratic elution (0.4 ml min^-1^; 10 % solvent A and 90 % solvent B, with A: 100 % 25 mM ammonium formate in ultra-pure water at pH 3.5 and B: 100 % acetonitrile) on a Waters Acquity BEH HILIC column (2.1 × 150 mm; 1.7 µm particle size) and a pre-column of the same material. Run time was 10 min and injection volume was 10 µl. The MS was equipped with an electrospray ionization source, which was operated in positive mode. MS parameters were optimized per repeated flow-injection analyses of TMA standards, as follows: drying gas flow 13 L/min; drying gas temperature 300 °C; nebulizer pressure 60 psi; capillary voltage 4,000 mV; fragmentor 120 V. Data were acquired in scan mode (10 - 200 *m*/*z*). A dilution series of TMA was used for quantification, with lowermost limits of detection and quantification equal to 0.01 ng and 0.1 ng TMA, respectively.

### Structure prediction and analysis

Protein structures of complexes were predicted using AlphaFold3 via the AlphaFold3 Server (Abramson et al., 2024) or for single monomeric proteins obtained from the AlphaFold Protein Structure Database (Varadi et al., 2024). Molecular graphics and analyses performed with UCSF Software ChimeraX (Meng et al., 2023) and Software Mole 2.5 for pore identification (Berka et al., 2017).

## Acknowledgments

We thank the AG Schmitz-Streit for fruitful discussions and critical comments. This work was supported by the German Research Foundation (DFG) priority program SPP2002 “Small Proteins in Prokaryotes, an Unexplored World” RSCHM1052/20-1 and 20-2, RSCHM1052/19-2 (to RAS), as well as DFG grant # 441217575 (to FJE).

## Conflict of interest

The authors declare that they have no conflicts of interest.

## Author contribution

RAS initiated the project. TH and RAS designed the experiments. TH performed the majority of the experiments. LH and CK contributed to the cloning. TH and FE performed the TMA transport assay using HPLC-MS. TH wrote the manuscript with RAS. TH and RAS interpreted and finalized the results. RAS supervised the research and provided resources and funding. All authors approved the submitted version.

## Supplemental tables and figures

**Fig. S1:**
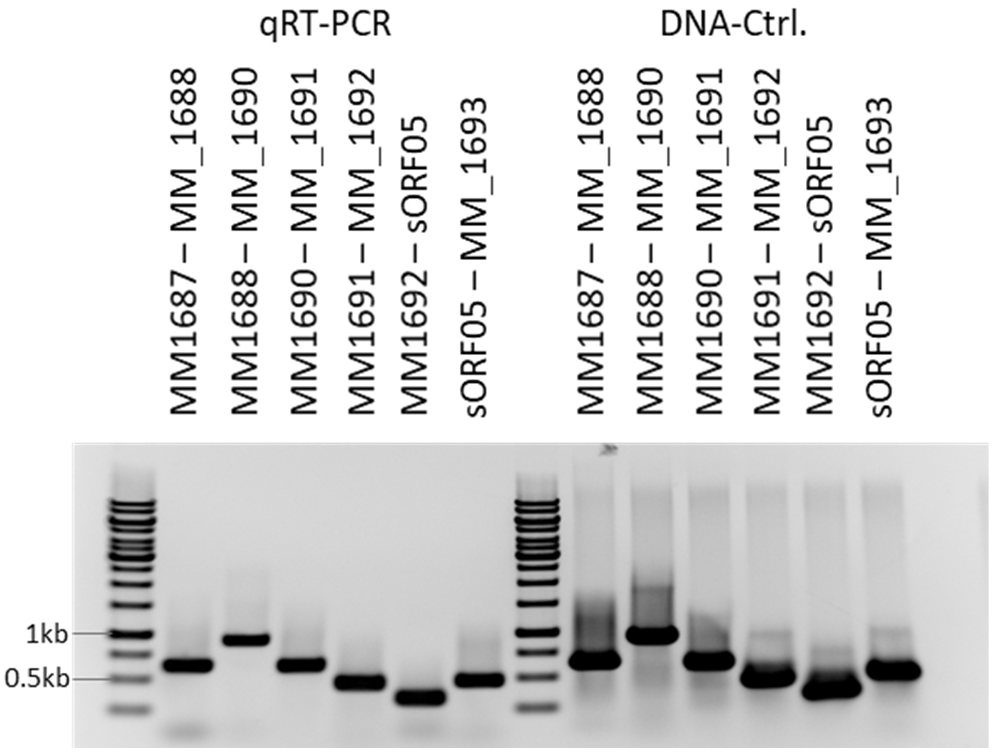
The genes MM_1687 – MM_1693 are organized in one operon. RT-PCR using primer pairs spanning from one gene to the next downstream gene were used to successfully amplify the target sequence regions, confirming that all seven genes are transcribed in one continuous transcript.

**Fig. S3:**
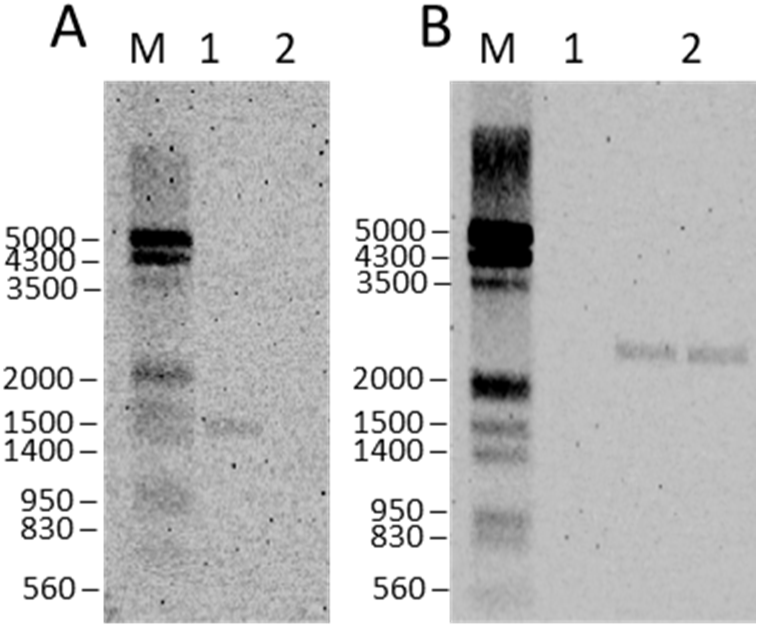
Southern Blot of the *M. mazei* WT (1) and the D*mttS* strain (2) with specific probes targeting the *mttS* gene (A) or the puromycin resistance cassette (B). The *mttS* is only present in the WT, while it has been exchanged with the pac in the D*mttS* strain confirming the successful genomic deletion of *mttS*.

**Table S1:**
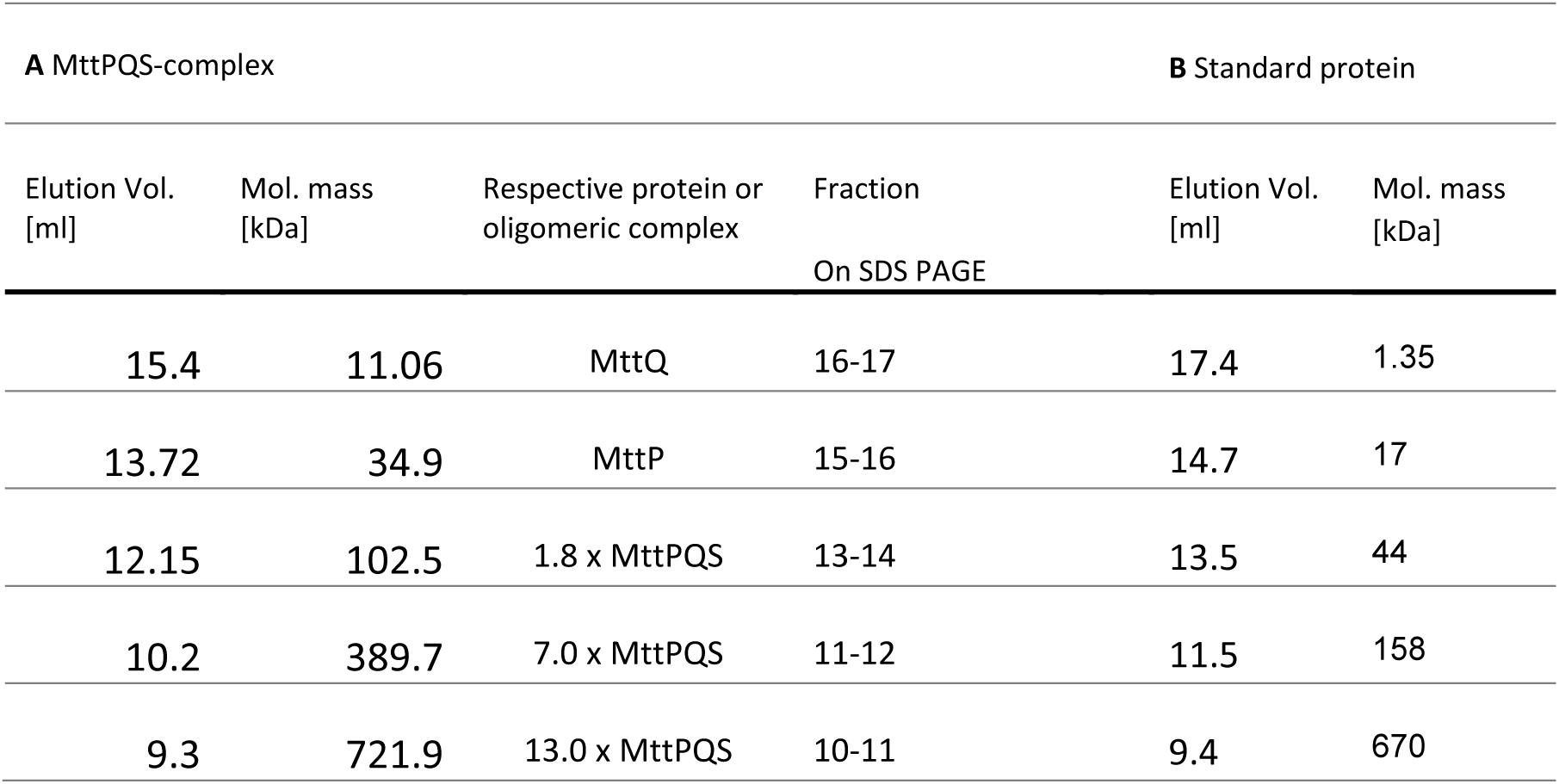
Calculated molecular masses corresponding to the SEC peaks. . For every local maximum of the size exclusion chromatogramm (A), the corresponding molecular mass was calculated by interpolation to the SEC standard (B). For every resulting molecular mass, the potential proteins are listed which fit the calculated mol. masses. Monomeric weight of MttQ 11 kDa, MttP 35 kDa, MttS 5.5 kDa, MttPQS protomer 55.5 kDa.

**T S2 Table:**
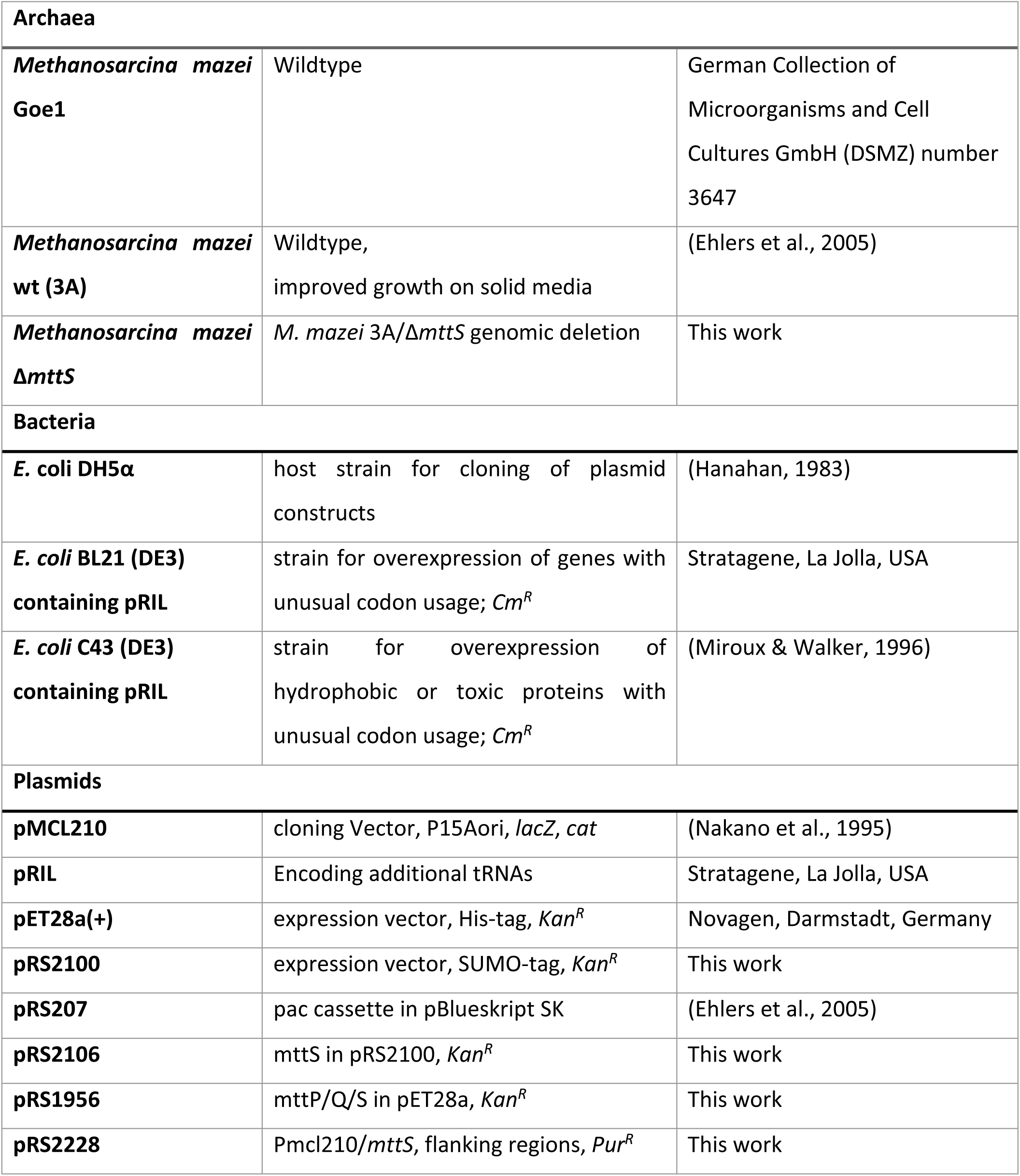
Used Strains and plasmids

**S3 Table:**
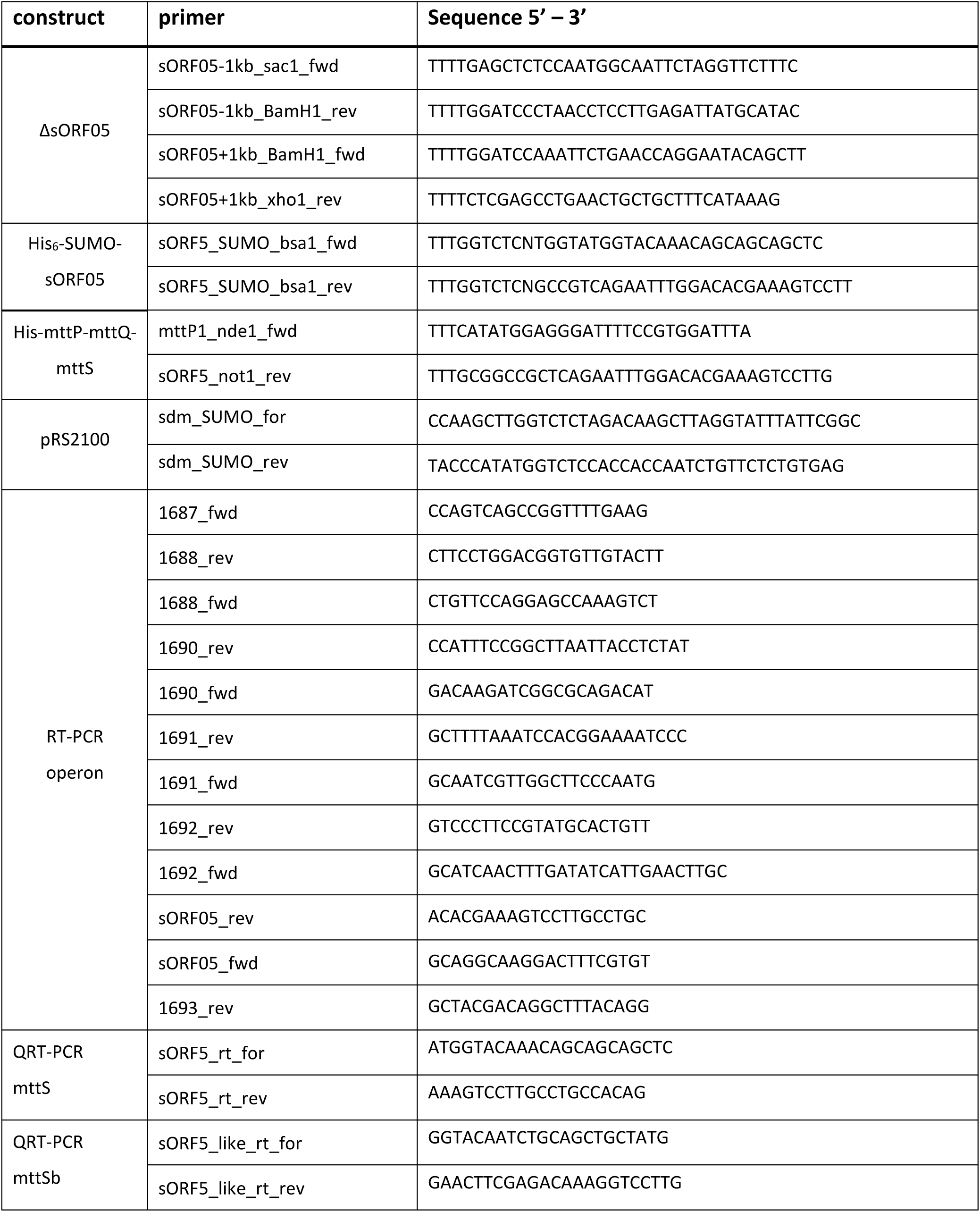
Used Oligonucleotides

## Notes

### Competing Interest Statement

The authors have declared no competing interest.

