## Supplementary material for "Small subunits MttS and MttQ of the MttP transporter regulate trimethylamine transport in *Methanosarcina mazei*": supplemetal figures and tables

### Supplemental tables and figures

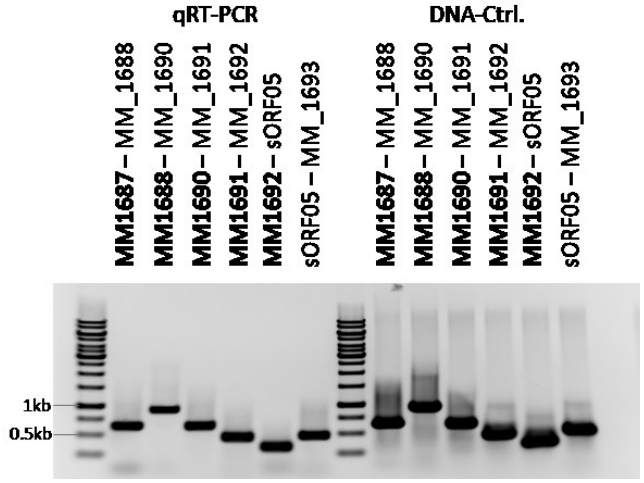


**Fig. S1: The genes MM_1687 – MM_1693 are organized in one operon.** RT-PCR using primer pairs spanning from one gene to the next downstream gene were used to successfully amplify the target sequence regions, confirming that all seven genes are transcribed in one continuous transcript.

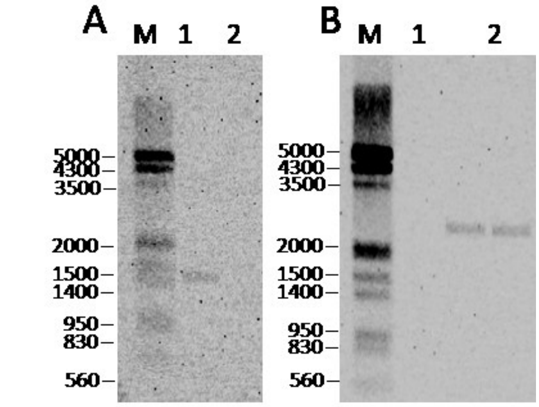


**Fig. S3:** Southern Blot of the *M. mazei* WT (1) and the D*mttS* strain (2) with specific probes targeting the *mttS* gene (A) or the puromycin resistance cassette (B). The *mttS* is only present in the WT, while it has been exchanged with the pac in the D*mttS* strain confirming the successful genomic deletion of *mttS*.

**Table S1: Calculated molecular masses corresponding to the SEC peaks**. For every local maximum of the size exclusion chromatogramm (A), the corresponding molecular mass was calculated by interpolation to the SEC standard (B). For every resulting molecular mass, the potential proteins are listed which fit the calculated mol. masses. Monomeric weight of MttQ 11 kDa, MttP 35 kDa, MttS 5.5 kDa, MttPQS protomer 55.5 kDa.

| **A** MttPQS-complex | | | | **B** Standard protein | |
|---|---|---|---|---|---|
| Elution Vol. [ml] | Mol. mass [kDa] | Respective protein or oligomeric complex | Fraction On SDS PAGE | Elution Vol. [ml] | Mol. mass [kDa] |
| 15.4 | 11.06 | MttQ | 16-17 | 17.4 | 1.35 |
| 13.72 | 34.9 | MttP | 15-16 | 14.7 | 17 |
| 12.15 | 102.5 | 1.8 x MttPQS | 13-14 | 13.5 | 44 |
| 10.2 | 389.7 | 7.0 x MttPQS | 11-12 | 11.5 | 158 |
| 9.3 | 721.9 | 13.0 x MttPQS | 10-11 | 9.4 | 670 |

S2 Table: Used Strains and plasmids

| | | |
|---|---|---|
| **Archaea** | | |
| ***Methanosarcina mazei* Goe1** | Wildtype | German Collection of Microorganisms and Cell Cultures GmbH (DSMZ) number 3647 |
| ***Methanosarcina mazei* wt (3A)** | Wildtype,<br>improved growth on solid media | (Ehlers et al., 2005) |
| ***Methanosarcina mazei* Δ*mttS*** | *M. mazei* 3A/Δ*mttS* genomic deletion | This work |
| **Bacteria** | | |
| ***E.* coli DH5α** | host strain for cloning of plasmid constructs | (Hanahan, 1983) |
| ***E. coli* BL21 (DE3) containing pRIL** | strain for overexpression of genes with unusual codon usage; $Cm^R$ | Stratagene, La Jolla, USA |
| ***E. coli* C43 (DE3) containing pRIL** | strain for overexpression of hydrophobic or toxic proteins with unusual codon usage; $Cm^R$ | (Miroux & Walker, 1996) |
| **Plasmids** | | |
| **pMCL210** | cloning Vector, P15Aori, *lacZ*, *cat* | (Nakano et al., 1995) |
| **pRIL** | Encoding additional tRNAs | Stratagene, La Jolla, USA |
| **pET28a(+)** | expression vector, His-tag, $Kan^R$ | Novagen, Darmstadt, Germany |
| **pRS2100** | expression vector, SUMO-tag, $Kan^R$ | This work |
| **pRS207** | pac cassette in pBlueskript SK | (Ehlers et al., 2005) |
| **pRS2106** | mttS in pRS2100, $Kan^R$ | This work |
| **pRS1956** | mttP/Q/S in pET28a, $Kan^R$ | This work |
| **pRS2228** | Pmcl210/*mttS*, flanking regions, $Pur^R$ | This work |

#### S3 Table: Used Oligonucleotides

| construct | primer | Sequence 5' – 3' |
|---|---|---|
| ΔsORF05 | sORF05-1kb_sac1_fwd | TTTTGAGCTCTCCAATGGCAATTCTAGGTTCTTTC |
| | sORF05-1kb_BamH1_rev | TTTTGGATCCCTAACCTCCTTGAGATTATGCATAC |
| | sORF05+1kb_BamH1_fwd | TTTTGGATCCAAATTCTGAACCAGGAATACAGCTT |
| | sORF05+1kb_xho1_rev | TTTTCTCGAGCCTGAACTGCTGCTTTCATAAAG |
| $His_6$-SUMO-sORF05 | sORF5_SUMO_bsa1_fwd | TTTGGTCTCNTGGTATGGTACAAACAGCAGCAGCTC |
| | sORF5_SUMO_bsa1_rev | TTTGGTCTCNGCCGTCAGAATTTGGACACGAAAGTCCTT |
| His-mttP-mttQ-mttS | mttP1_nde1_fwd | TTTCATATGGAGGGATTTTCCGTGGATTTA |
| | sORF5_not1_rev | TTTGCGGCCGCTCAGAATTTGGACACGAAAGTCCTTG |
| pRS2100 | sdm_SUMO_for | CCAAGCTTGGTCTCTAGACAAGCTTAGGTATTTATTCGGC |
| | sdm_SUMO_rev | TACCCATATGGTCTCCACCACCAATCTGTTCTCTGTGAG |
| RT-PCR operon | 1687_fwd | CCAGTCAGCCGGTTTTGAAG |
| | 1688_rev | CTTCCTGGACGGTGTTGTACTT |
| | 1688_fwd | CTGTTCCAGGAGCCAAAGTCT |
| | 1690_rev | CCATTTCCGGCTTAATTACCTCTAT |
| | 1690_fwd | GACAAGATCGGCGCAGACAT |
| | 1691_rev | GCTTTTAAATCCACGGAAAATCCC |
| | 1691_fwd | GCAATCGTTGGCTTCCCAATG |
| | 1692_rev | GTCCCTTCCGTATGCACTGTT |
| | 1692_fwd | GCATCAACTTTGATATCATTGAACTTGC |
| | sORF05_rev | ACACGAAAGTCCTTGCCTGC |
| | sORF05_fwd | GCAGGCAAGGACTTTCGTGT |
| | 1693_rev | GCTACGACAGGCTTTACAGG |
| QRT-PCR mttS | sORF5_rt_for | ATGGTACAAACAGCAGCAGCTC |
| | sORF5_rt_rev | AAAGTCCTTGCCTGCCACAG |
| QRT-PCR mttSb | sORF5_like_rt_for | GGTACAATCTGCAGCTGCTATG |
| | sORF5_like_rt_rev | GAACTTCGAGACAAAGGTCCTTG |

Ehlers, C., Weidenbach, K., Veit, K., Deppenmeier, U., Metcalf, W. W., & Schmitz, R. A. (2005). Development of genetic methods and construction of a chromosomal glnK 1 mutant in Methanosarcina mazei strain Gö1. *Molecular Genetics and Genomics*, *273*(4), 290–298. https://doi.org/10.1007/s00438-005-1128-7

Hanahan, D. (1983). Studies on transformation of Escherichia coli with plasmids. *Journal of Molecular Biology*, *166*(4), 557–580. https://doi.org/10.1016/S0022-2836(83)80284-8

Miroux, B., & Walker, J. E. (1996). Over-production of Proteins in Escherichia coli: Mutant Hosts that Allow Synthesis of some Membrane Proteins and Globular Proteins at High Levels. In *J. Mol. Biol* (Vol. 260).

Nakano, Y., Yoshida, Y., Yamashita, Y., & Koga, T. (1995). Construction of a series of pACYC-derived plasmid vectors. *Gene*, *162*(1), 157–158. https://doi.org/10.1016/0378-1119(95)00320-6
